# Sex differences in the nasal microbiome of healthy young adults

**DOI:** 10.1101/2022.05.23.493011

**Authors:** Yanmei Ju, Zhe Zhang, Mingliang Liu, Shutian Lin, Qiang Sun, Zewei Song, Weiting Liang, Xin Tong, Zhuye Jie, Haorong Lu, Kaiye Cai, Peishan Chen, Xin Jin, Xun Xu, Huanming Yang, Jian Wang, Yong Hou, Liang Xiao, Huijue Jia, Tao Zhang, Ruijin Guo

## Abstract

**Respiratory diseases impose an immense health burden worldwide. Epidemiological studies have revealed extensive disparities in the incidence and severity of respiratory tract infections (RTIs) between males and females**. It is recently hypothesized that there might also be a nasal microbiome axis contributing to the observed sex disparities, but without evidence. In this work, we study the **nasal microbiome of healthy young adults** in, as of today, **the largest cohort** based on deep shot-gun metagenomic sequencing. We mainly focus on the bacteriome, but also integrate the mycobiome to get a more holistic perspective. **De novo assembly** is performed to catalog the nasal bacterial colonizers/residents, which also **identify and therefore account for uncharacterized components** of the community. The bacteriome is then profiled based on the non-redundant metagenome-assembled genomes **(MAGs) catalog** constructed therefrom. Unsupervised clustering reveals clearly separable structural patterns in the nasal microbiome between the two sexes. Following this link, we **systematically evaluate sex differences for the first time and reveal extensive sex-specific features in the nasal microbiome composition. More importantly, through network analyses, we capture markedly higher ecological stability and antagonistic potentials in the nasal microbiome of females than that of males. The analysis of the keystone bacteria of the communities reveal that the sex-dependent evolutionary characteristics might have contributed to this difference**.

**Highlights:** The non-redundant nasal bacterial MAGs catalog constructed from ultra-deeply sequenced metagenomic data provides a valuable resource.

Integrating nasal bacteriome and mycobiome data provides a more holistic perspective for the understudied human nasal microbiome.

Unsupervised clustering helps uncover extensive sex differences in the nasal microbiome compositions.

Network analyses capture markedly higher ecological stability and antagonistic potentials in the nasal microbiome of females than that of males.

Sex-dependent genetic evolutionary forces play a role in the shaping of keystones in the nasal microbial community.

## Introduction

Respiratory diseases impose an immense health burden worldwide, affecting billions of people’s lives and accounting for over 10% of all disability-adjusted life-years (DALY) as of 2019 according to the Global Burden of Diseases (GBD) study (Ferkol and Schraufnagel, 2014; GBD 2019 Diseases and Injuries Collaborators, 2020; Jin et al., 2021; Viegi et al., 2020), let alone the catastrophic impact of the COVID-19 pandemic. Sex is a significant factor in many diseases. Epidemiological studies have revealed extensive disparities in the incidence and severity of respiratory tract infections (RTIs) between males and females. Males are generally more commonly and severely affected by most RTIs than females across all age groups (Falagas et al., 2007; Jacobsen and Klein, 2021; Jin et al., 2021). A greater mortality rate for males was also observed in COVID-19

(Klein et al., 2020; Scully et al., 2020; Vahidy et al., 2021). Sex-specific differences in immunity mediated by sex chromosome complement, genes and sex hormones can play important roles in the observed disparity (Jacobsen and Klein, 2021; Scully et al., 2020; Vahidy et al., 2021). Nevertheless, the mechanism remains unclear. The nasal microbiome has been implicated in different respiratory diseases (Fazlollahi et al., 2018; Gan et al., 2021; Kumpitsch et al., 2019; Mahdavinia et al., 2016; Ramakrishnan and Frank, 2018; Rhee et al., 2021; B. G. Wu et al., 2019). It is recently proposed that there might also be a nasal microbiome axis contributing to the observed sex disparities (Shah, 2021).

The nasal bacterial community is characterized by a high prevalence of *Corynebacterium spp*., *Propionibacterium spp*. and *Staphylococcus spp*., with most components belonging to phyla Actinobacteria, Firmicutes and Proteobacteria (Human Microbiome Project Consortium, 2012; C. M. Liu et al., 2015). In addition to bacterial colonizers, the nasal cavity also harbors a mycobiota (Jung et al., 2015), as well as the presence of viruses. However, the nasal microbiome studies are hitherto still limited to small sample sizes or 16S rRNA gene-based sequencing (Wenkui Dai et al., 2019; de Steenhuijsen Piters et al., 2020; Earl et al., 2018; Kaul et al., 2020; C. M. Liu et al., 2015; Yan et al., 2013). Sex differences in the nasal microbiome have never been systematically evaluated. Liu *et al*. identified seven community state types (CSTs) of the nasal bacterial community in a cohort of 86 twin pairs above 50 years old, and found no significant difference in the CST distribution between the two sexes despite higher microbial loads in the nasal cavity of males. This is not surprising considering that even in the most researched gut microbiome, sex differences only came to light very recently through large-scale population studies (la Cuesta-Zuluaga et al., 2019; Sinha et al., 2018; X. Zhang et al., 2021). In a well-designed large cohort, which nicely limited variability in potential confounding factors, such as genetic background, geographic residence, and diet habits etc., Zhang et al. demonstrated sex-specific aging trajectories of the gut microbiome. They showed that the differences were especially evident between age-matched male adults and premenopausal female adults (approximately below ∼50 years old), and gradually diminished after 50 years of age.

It is increasingly recognized that the nasal microbiome might function as a gatekeeper in respiratory health (Wenfang Dai et al., 2018; Man et al., 2017). The nasal cavity is featured by limited nutrients and adhesion surfaces (Krismer et al., 2014), and represents a major reservoir for opportunistic pathogens, such as *Staphylococcus aureus, Streptococcus pneumoniae*, and *Haemophilus influenzae* (Clark, 2020). The microbes in this niche are hence in constant competition, and sometimes form cooperative relation, to gain self-fitness (Brugger et al., 2020; De Boeck et al., 2021; Zipperer et al., 2016). Extensive antimicrobial substance productions have been identified in nasal microbes, which can be potential mediators of the interactions (De Boeck et al., 2021; Donia et al., 2014; Donia and Fischbach, 2015; Iwase et al., 2010; Janek et al., 2016; Zipperer et al., 2016). The competitive (antagonistic) and cooperative (synergistic) interactions influence both the initial colonization of pathogens and the thereafter dynamics. Network-based approaches have been proved to be helpful in deciphering complex interactions and are increasingly applied in the microbial field. Understanding the nature of microbial co-occurrence and correlation patterns within and cross domains may provide insights into the ecological systems as well as related human diseases. Through network-based analyses, researchers studying bronchiectasis exacerbations found that patients of different exacerbation risks featured distinct microbial interaction networks (Mac Aogáin et al., 2021). Instead of the implicated single pathobiont *Pseudomonas*, it is the interaction network that is associated with the exacerbation risk. While cross-domain interactions are rarely explored, Tipton et al. recently showed that compared to single domain networks, bacteria-fungi combined networks had higher overall connectivity and increased attack robustness (Tipton et al., 2018). More importantly, network analyses can help elucidate and prioritize the keystones of a community, which may not be the species dominant in abundance, and sometimes can even be unknown “microbial dark matter” (Banerjee et al., 2018; Zamkovaya et al., 2020).

In this work, we study the nasal microbiome of healthy young adults in a so far largest cohort based on deep shotgun metagenomic sequencing. We mainly focus on the bacteriome, but also integrate the mycobiome to get a more holistic perspective. De novo assembly is performed to catalog the nasal bacterial colonizers/residents, which also identify and therefore account for uncharacterized components of the community. The bacteriome is then profiled based on the non-redundant metagenome-assembled genomes (MAGs) catalog constructed therefrom. Unsupervised clustering reveals clearly separable patterns between the two sexes, implying a distinct structure of the nasal microbiome between males and females. Following this link, we systematically evaluate sex differences for the first time and reveal extensive sex-specific features in the nasal microbiome composition. More importantly, through network analyses, we capture markedly higher stability and antagonistic potentials in the nasal microbiome of females than that of males, in the shaping of which the sex-dependent evolutionary characteristics might have played a role as revealed by the keystone bacteria of the communities.

## Results

### Characterizing the nasal bacteriome and mycobiome

To characterize the nasal microbiome of healthy young adults, we performed deep shotgun metagenomic sequencing on 1,593 anterior nares samples from the 4D-SZ cohort (C. Chen et al., 2021; Jie et al., 2021a; 2021b; X. Liu et al., 2022; Zhu et al., 2021) (Table S1). In total, 128.21 terabases raw data were generated with an average of 80.48 gigabases for each sample (Table S2). A single sample assembly, and single sample binning strategy (see Methods) was employed to reconstruct genomes from the ultra-deeply sequenced metagenomic data. A total of 4,197 metagenome-assembled genomes (MAGs) were assembled at a threshold for quality control of >50% completeness and <= 10% contamination. To compile a non-redundant MAGs catalog, we performed de-replication with 99% of the average nucleotide identity (ANI). At the end, a catalog of 974 non-redundant MAGs for human nasal associated bacteria were retained, which included 718 high-quality (completeness > 90% & contamination < 5%) and 256 medium quality (completeness > 50% & contamination < 10%) ones (Figure 1a; Figure S1). 16S rRNA genes had been detected in about 45% of the 974 MAGs (Figure 1a). To explore the taxonomic coverage of this catalog, we classified the MAGs according to 95% average nucleotide identity. Overall, we obtained 232 species from 13 known phyla, with 150 annotated to known genomes in the GTDB database, and the other 82 as newly identified (unknown) (Figure. 1b). The unknown species spanned over all of the 13 phyla, with the largest number from Bacteroidota. For four phyla, including Fusobacteriota, Eremiobacterota, Deinococcota and Bdellovibrionota, only unknown species were discovered. This suggests that the habitat of the nasal cavity featured drastically distinct characteristics from other habitats where the species of these phyla are often identified, e.g. *Fusobacterium nucleatum* of phylum Fusobacteriota is often detected in oral and fecal samples. Additionally, we identified six novel genera from phylum Proteobacteria, Bacteroidota and Firmicutes_A, and one novel family from phylum Eremiobacterota, which cannot be assigned to any known taxa of the respective level at the level-specific phylogenetic distance cut-offs (Table S3). Notably, our data also improved the genome completeness of a singleton taxon, namely *QFNR01 sp003248485* (90.42% completeness, compared with 75.15% completeness in GTDB; Table S3). Overall, the majority of the MAGs in the catalog belonged to Actinobacteriota, Proteobacteria and Firmicutes, which is typical for the human nasal microbiome (Clark, 2020; Dimitri-Pinheiro et al., 2020; Human Microbiome Project Consortium, 2012).

**Fig. 1.**
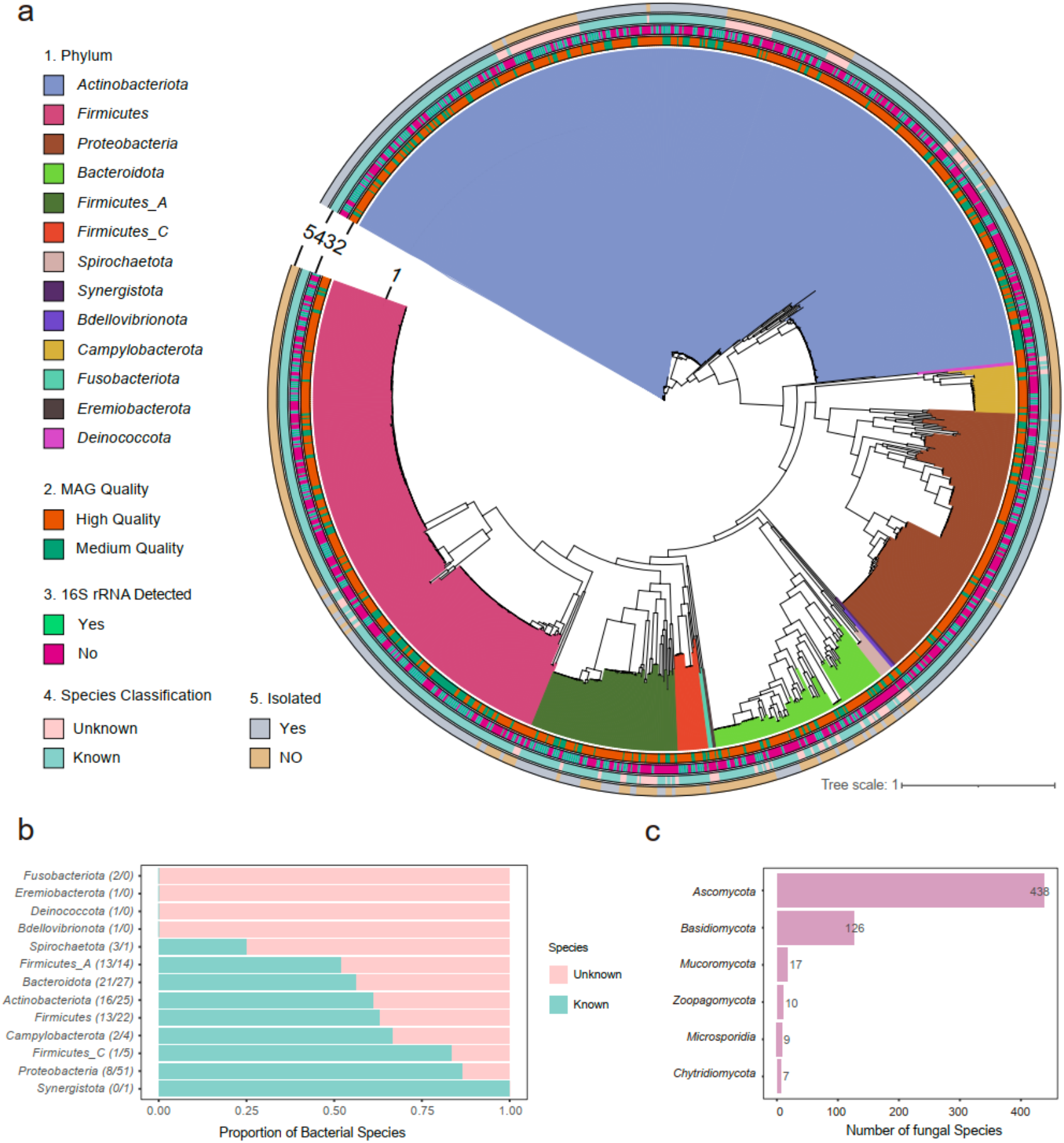
Overall representation of the microbes in anterior nares of healthy young adults. **a**, Phylogeny of 974 non-redundant bacterial MAGs (metagenome-assembled genomes) detected in anterior nares. It constituted five layers representing respectively: 1 for phylum, 2 for MAGs quality, 3 for if 16s rRNA detected, 4 for if classified in the species level and 5 for if isolated as depicted in the GTDB database. **b**, Proportion of unknown and known bacterial species in each phylum with the absolute number indicated in the brackets respectively. **c**, Number of fungal species in each phylum.

Natural products of human microbiota are increasingly recognized as important mediators for a variety of microbe-host and microbe-microbe interactions, which in turn can be explored for potential pharmaceutical applications (Donia and Fischbach, 2015; Milshteyn et al., 2018; Sugimoto et al., 2019). As an example, a nasal isolate of *Staphylococcus lugdunensis* has recently been shown to produce a novel antibiotic, lugdunin, a non-ribosomally synthesized bioactive natural product, which is bactericidal against major human pathogens and prohibits the colonization of *S. aureus* in the nasal cavity (Zipperer et al., 2016). We therefore screened for the presence of secondary metabolites biosynthetic gene clusters (BGCs) encoded within the 974 non-redundant MAGs using antiSMASH (Blin et al., 2013; 2018; Medema et al., 2011) (Table S3b). In total, we detected 2,921 BGCs, which were primarily inferred as synthesized terpenes, nonribosomal peptides (NRPs), types I polyketide synthases (PKSs), siderophores and other unspecified ribosomally synthesised and post-translationally modified peptide products (RiPPs). Notably, 514 of them were screened from MAGs newly identified from the nasal microbiome in this cohort (Figure S2b). In addition, 1,975 (67.2%) of the detected gene clusters were novel clusters, most of which were from Actinobacteriota, Firmicutes and Proteobacteria (Figure S2c). These data, in particular the high number and proportion of novel clusters, suggests that the nasal microbiota may serve as a rich reservoir for new antibiotics or other pharmaceuticals.

To profile the nasal microbiome composition, the metagenome data were first mapped to the constructed non-redundant nasal MAGs catalog and generated the bacterial profile for each sample. The most abundant species were mainly from genera *Corynebacterium, Staphylococcus, Moraxella, Cutibacterium, Dolosigranulum*, and *Lawsonella*. Accumulative abundance analysis showed that top-ranked 17 species accounted for over 90% of the overall composition, suggesting that the nasal bacterial community was dominated by a few taxa (Figure S3). To characterize the mycobiome composition, we aligned our high-quality cleaned metagenome data to a manually curated database with Kraken. In total, we identified 607 fungal species in this cohort with the most from phyla Ascomycota and Basidiomycota (Figure 1c). While the bacterial community was dominated by several species, the mycobiome was more evenly distributed, taking over 50 species to account for ∼90% of the overall fungal mycobiome composition. *Aspergillus spp*. and *Malasseziaceae spp*., among others, are the most abundant fungi in the nasal cavity of this cohort (Figure S3).

### Unsupervised clustering helps uncover sex differences in the nasal microbiome composition

To gain a holistic perspective of the microbial structure, we integrated the bacterial and fungal community profile with a weighted similarity network fusion approach. Unsupervised clustering of the resultant similarity matrix classified the cohort into three clusters (Figure 2a; see Methods). Permutational multivariate analysis of variance (PERMANOVA) on bray dissimilarity showed that these clusters explained over 18% of the variance in the composition between any two or among all three clusters (Figure 2b). In particular clusters 2 and 3 contained samples almost exclusively from single sex, i.e. male and female, respectively. Even in cluster 1 the similarity matrix featured two separable patterns corresponding to two sexes, suggesting distinctive structures in the nasal microbiome composition between males and females.

**Figure 2.**
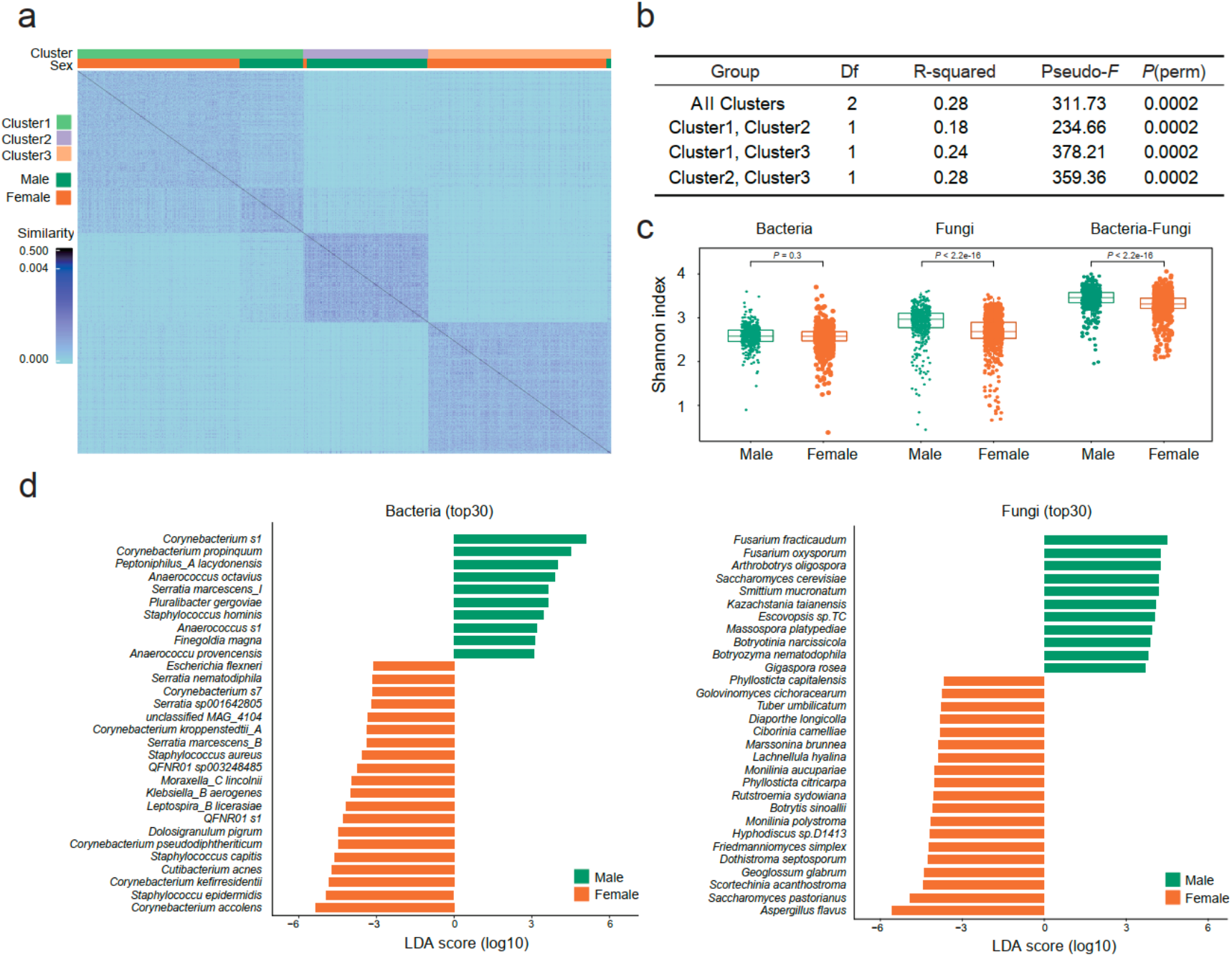
Unsupervised clustering and sex differences in the nasal microbiome composition. **a**, Heatmap illustrating WSNF similarity scores stratified by unsupervised clustering through bacterial and fungal datasets, with cluster and sex information indicated by the bars on the top. **b**, Permanova analysis (permutational multivariate analysis of variance using distance matrices) results demonstrating significant variation of the nasal microbiome among the observed clusters. **c**, Comparison of α-diversity (Shannon index) between males (green) and females (brown) for bacteria (*P*=0.3), fungi (*P*<2.2×10^−16^) and bacteria-fungi integrated (*P*<2.2×10^−16^) respectively through Wilcoxon test. **d**, Comparison of LDA effect size (LEfSe) between males (green) and females (brown) illustrating discriminative taxa of bacteria (left) and fungi (right). Only the top 30 discriminative taxa by LDA score were shown.

Following this link that unsupervised clustering uncovered, we systematically evaluated the sex differences in the nasal microbiome composition. PERMANOVA confirmed that sex was a significant covariant for the nasal bacteriome, mycobiome as well as the bacteria-fungi integrated microbiome (Table S4). In the bacteria-fungi integrated profile we observed a significantly higher Shannon diversity in males than in females (Figure 2c; Table S5). However, this observation was mainly attributed to the mycobiome. No significant difference in the Shannon diversity of the bacteriome was detected. For individual microbial taxon, we performed linear discriminative analysis (LDA) and identified considerable significant associations between the relative abundances and sex. Specifically, at the species level 59 bacteria and 148 fungi, and at the genus level 24 bacteria and 23 fungi, were significantly different in abundance between males and females (Figure 2d; Table S6). Interestingly, the taxa number enriched in females almost doubled that in males. Notably, *Staphylococcus aureus*, a commonly known opportunistic pathogen in the nasal cavity, was not only more prevalent but also significantly more enriched in females. Meanwhile, *Corynebacterium accolens, Corynebacterium pseudodiphtheriticum*, and *Dolosigranulum pigrum*, for which cooperative or competitive relationships with *S. aureus* have been identified in former studies via association and experimental validation (Brugger et al., 2020; De Boeck et al., 2021; Krismer et al., 2017; Yan et al., 2013), as the most abundant species among others in this cohort, were also significantly more abundant in females. Additionally, *Lactobacillus spp*., typically found in the female vagina, has been recently reported to have a niche in the human nose and may exert a beneficial effect (De Boeck et al., 2020). In our cohort, *Lactobacillus spp*. were also detected, with females characterized by a higher relative abundance of *L. iners* and *L. crispatus* than males.

### Network analyses capture markedly higher ecological stability and antagonistic potentials in the nasal microbiome of females than that of males

Having uncovered extensive sex differences in the microbial composition, next we aimed to determine if the nasal microbiome featured different ecological relationship characteristics between males and females. To characterize the microbial interactions within each sex, we employed an integrated approach combining COAT (composition-adjusted thresholding), HUGE (High-dimensional Undirected Graph Estimation), MI (mutual information) and Bray-Curtis dissimilarity to construct the co-occurrence networks (Figure 3a; see Methods). The inferred interactions between microbes, as nodes in the graphs, were represented by signed edges in the network, with positive for cooperative/synergistic relation and negative for competitive/antagonistic relation. The total number of interactions (edges) was very close between the two sexes with a marginally higher number of negative interactions in females. Splitting the entire network into three sub-networks, i.e. within the bacteria domain, within the fungi domain and cross bacteria-fungi domain, revealed that the cross-domain sub-network accounted for over half of the negative interactions (Figure 3b; Table S7).

**Fig3.**
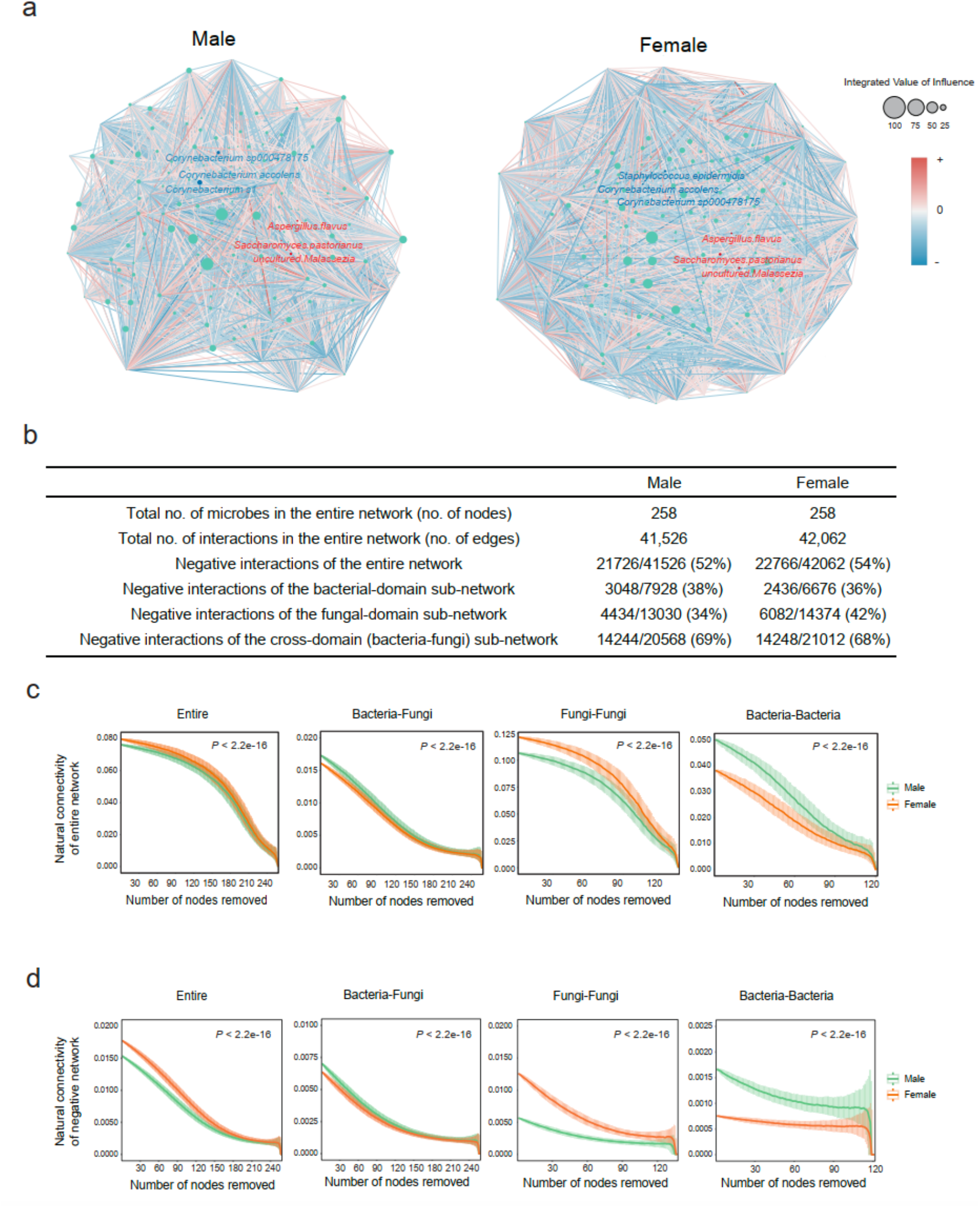
Network characterization of the nasal microbiome for males and females. **a**, Nasal microbial interaction network of males and females. Node size represents the integrated value of influence (IVI) for each taxon. Red and blue lines indicate positive and negative interactions respectively. The top 3 bacteria and fungi by relative abundance are annotated with blue and red fonts respectively. **b**, Summary table of network characteristics of males and females **c-d**, Attack robustness of each network of total interaction (**c**) and negative interaction (**d**) for males (green) and females (brown) as measured by natural connectivity. Line and box reflect median and IQRs.

The functioning of complex networks largely relies on their robustness (Matchado et al., 2021), a better understanding of which can provide valuable insights into RTI susceptibility and pathologies. We thus adopted a sensitive and reliable measure, namely natural connectivity (Peng and J. Wu, 2016; J. Wu et al., 2010; X.-K. Zhang et al., 2013), to quantify the stability of the inferred networks. To simulate the influence of microbes’ loss on the network, we performed random attacks and assessed the stability of the remaining network (hamster et al., 2019) (see Methods). Intriguingly, the network robustness was much higher for females than for males. While the human nasal microbiome is increasingly regarded as a gatekeeper of respiratory health, opportunistic pathogens do often present even in healthy individuals. Thus, the negative/antagonistic interactions are of particular interest. When only considering the negative interactions, we observed that much higher robustness for females still held. Significant separations of the natural connectivity plot were observed between the networks of males and females, for both the entire network and the negative network (P-value < 2.2e-16; Figure 3c and 3d). Higher overall natural connectivity for females largely remained until over half of the species were removed, further confirming that females characterized a more stable network with more intensive interactions and higher antagonistic potentials which may provide stronger resistance against opportunistic pathogens. Additionally, comparison of the random attack results between the cross-domain network and within domain networks demonstrated that the fungal mycobiome made a great contribution to the network robustness. Interestingly, a recent study also suggested that fungi played a stabilizing role in the lung and skin microbial ecosystems (Tipton et al., 2018).

### Sex-dependent genetic evolutionary forces in the shaping of keystones in the nasal microbial community

Network analysis can be a powerful tool for inferring keystone taxa of the microbial communities (Banerjee et al., 2018; Layeghifard et al., 2016; Matchado et al., 2021; Zamkovaya et al., 2020). To this end, we adopted a novel influential node detection method, integrated value of influence (IVI), which captures all topological dimensions of the networks, to assess the importance of individual taxon of the community (Salavaty et al., 2020). Notably, the IVIs of most taxa derived from the entire networks of males and females were considerably different (Table S8), indicating different levels of importance of the respective taxa potentially eliciting in the microbial community of each sex. Moreover, IVI only weakly correlated with relative abundance (Figure S4), suggesting that the most abundant taxa may not necessarily exert the strongest influences in the community (Banerjee et al., 2018; Zamkovaya et al., 2020). Keystone microbes represent the ones contributing the most to the robustness of the community. With a permutational approach (see Methods) we derived the keystone sets for males and females, which included 13 and 10 taxa respectively (Figure 4a). Intriguingly, the keystone sets for males and females both contained taxa from bacterial and fungal domains, but with completely different specific components and remarkably different IVIs between the two sexes for each keystone.

**Fig 4.**
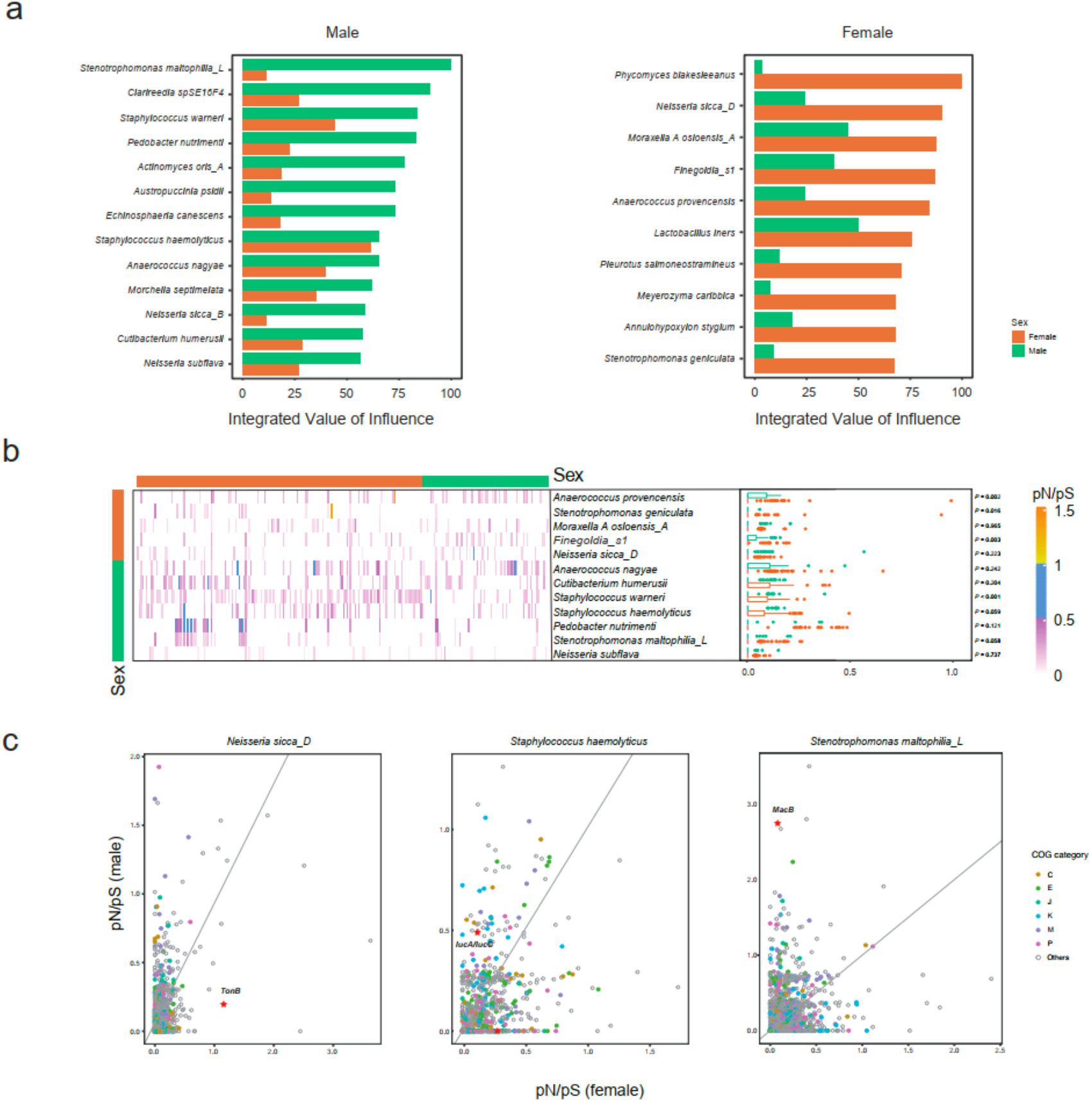
Characteristics of the keystone taxa identified in male and female nasal microbial interaction networks. **a**, The integrated value of influence (IVI) of the keystone taxa of males (left) and females (right). Green and brown bars represent the IVI of the respective taxa in the male and female networks respectively. **b**, The pN/pS ratio of keystone bacteria for individuals shown by heatmap and boxplot. Green and brown represent male and female respectively (vertical bar: keystone belongs to male or female network; horizontal bar: male and female individuals; boxplot: pN/pS ratios for male and female individuals). **c**, The pN/pS ratio for 3 bacterial keystones in gene levels of male (y-axis) and female (x-axis) participants with COG category. The stars represent the genes which have been described in detail in the main text.

Evolution is important for ecological dynamics in bacterial communities. To illuminate the genetic evolutionary characteristics of the keystones for each sex, we evaluated the selection of environmental pressures for the keystone bacteria by estimating the pN/pS ratio within each genome for each sample (Garud and Pollard, 2022; Schloissnig et al., 2013). The results showed that the pN/pS ratios varied among different species, but were mostly below one for both males and females (Figure 4b). This suggested that the evolution of the keystone bacteria was largely predominated by long-term purifying selection. On the other hand, the pN/pS ratios differed significantly between males and females in some of the keystone bacteria, including male-specific keystone *Staphylococcus warneri* and female-specific keystone *Anaerococcus provencensis, Stenotrophomonas geniculata*, and *Finegoldia s1* (Figure 4b). This can potentially be in relation to sex-specific evolutionary constraints confronted by the microbes in the nasal cavity of males and females, such as different levels of immunoinflammatory characteristics.

On the gene level, however, we observed considerable deviations in pN/pS ratios of the same keystone taxa between males and females, indicating sex-dependent selective pressures and genetic adaptations (Figure 4c, Figure S5; Table S9). For instance, the nasal cavity is noted for limited resources available, such as iron limitation (Krismer et al., 2014; Kumpitsch et al., 2019; Stubbendieck et al., 2019). Notably, the 974 nasal bacterial MAGs encoded remarkably more siderophores (Figure S2b), one of the main mechanisms for bacterial iron sequestering, compared to that detected in the large collection of human gut bacterial MAGs derived from over 10,000 samples (Almeida et al., 2019). Although *Neisseria sicca*, a common nasopharyngeal commensal, does not encode siderophores (Marri et al., 2010), we found the female keystone *N. sicca_D* underwent positive adaptation in genes encoding TonB-dependant siderophore receptors (mean pN/pS ratio of 1.168) in females, with which the bacteria can exploit siderophores produced by other members of the community for iron sequestering. In contrast, in males it was subjected to purifying selection in these genes with a mean pN/pS ratio of 0.195. In a male keystone bacterium, *Staphylococcus haemolyticus*, we also observed that genes encoding IucA/IucC family siderophore biosynthesis protein showed relaxed purifying selection in males (mean pN/pS ratio of 0.496) but tight purified selection in females (mean pN/pS ratio of 0.116). Gene *yfmC in S. haemolyticus*, which encodes Fe(^3+^)-citrate-binding protein involved in iron transport, was purged in males (mean pN/pS ration of 0) whereas showed tight purified selection (mean pN/pS ratio of 0.259) in females. Antibiotics represent another major category of stresses for bacteria, for which resistance evolves over time. As a global emerging multidrug-resistant organism, *Stenotrophomonas maltophilia* has been most commonly associated with respiratory infections in humans (Brooke, 2012) and isolated predominantly in elderly males of hospitalized lower RTI patients (Chawla et al., 2014). Like the other MacA-MacB-TolC tripartite efflux pumps, *S. maltophilia* MacB has been previously revealed to drive resistance to a variety of antibiotics, such as macrolides, aminoglycosides and polymyxins, in concert with MacA adaptor protein and TolC outer membrane exit duct (Crow et al., 2017; Koronakis, 2018; Lin et al., 2014). Interestingly, we found in *S. maltophilia_L* the gene coding for MacB exhibited strong positive selection in males, but tight negative selection in females (mean pN/pS ratio: 2.73 vs. 0.08). Together, the keystone bacteria exhibited highly sex-specific genetic evolutionary characteristics in niche-specific or sex-biased stress-related functional units, which was in close relation to their role in the respective network of each sex. This suggests that the genetic evolutionary forces might have played a role in the shaping of the keystones of the nasal microbial community of each sex. The effect might even be mutual, such that interactions of the keystones spurred evolution which in turn reinforced their role as keystone, or the other way around.

## Discussion

With advances in sequencing technologies, microbial research is no more restricted to cultivation. Great efforts have since been made to characterize the human microbiome. However, most of the studies rely on 16S rDNA amplicon-based or gene-centric microbial community characterization, which is heavily skewed by microbes that are easily cultivatable or the most researched habitats’ residents, such as the human gut microbiome (Bolyen et al., 2019; Callahan et al., 2016; MetaHIT Consortium et al., 2014; Segata et al., 2012; Woyke, 2019; Xie et al., 2016). Recently, genome-resolved metagenomics through de novo assembly has transformed our understanding of the microbiome composition, which can meanwhile provide valuable knowledge of individual species for deciphering their biological roles. The human microbiome has a strong niche specialization both within and among individuals (Human Microbiome Project Consortium, 2012). Large reference genome catalogs have been constructed for the human gut and oral microbiome, and massively expanded the known species repertoire of the respective habitats (Almeida et al., 2019; 2021; Nayfach et al., 2019; Pasolli et al., 2019; Zhu et al., 2021). Here in this work, we leveraged ultra-deeply sequenced metagenome data from a large cohort of healthy young adults and constructed a non-redundant nasal associated bacterial MAGs catalog. It represents the first endeavor in cataloging the human nasal microbial reference genome, and makes a great contribution to the global effort for characterizing the human microbiome. The catalog provides a valuable resource for profiling the nasal microbiome and developing new antibiotics or other pharmaceuticals in future studies. Meanwhile, it makes it possible for uncovering potentially important unknown taxa in this ecosystem.

Respiratory health is of vital importance for human beings. The COVID-19 pandemic has made it unprecedentedly clear. Sex biases have been widely noted in different types of respiratory diseases, COVID-19 included as well. Heightened immunity in females renders them generally less affected by infections, but more prone to autoimmunity diseases (Jacobsen and Klein, 2021; Scully et al., 2020; Vahidy et al., 2021). Sex hormones and chromosomes can also play important roles. However the underlying mechanisms remain unclear. Recently it has been argued that the nasal microbiome might also play a role in the observed disparities between males and females, but unfortunately lacked support and evidence (Shah, 2021). In this work, unsupervised clustering of the nasal microbiota revealed clearly separable patterns between males and females. This led us to further systematically evaluate the sex differences in this community and uncovered extensive sex-specific features. Females harbored higher abundances of more taxa, including the commonly known respiratory tract opportunistic pathogen, *Staphylococcus aureus*, and several species formerly identified associating with it. Intriguingly, the interaction networks of females also featured higher robustness and stronger antagonistic interaction potentials than males. The connection of such characteristics with lower susceptibility and severity of RTIs in females compared to males warrants further investigation. While bacterial interaction networks are widely studied and cross-domain interactions are rarely explored, our work integrated the bacteriome and mycobiome and gained a more holistic perspective of the community. Our results suggested that the mycobiome might play an important stabilizing role, in an echo of a former study (Tipton et al., 2018). Through network analysis, we identified sex-specific keystone microbes, which also included formerly unknown taxa, demonstrating the power and necessity of cataloging the community through de novo assembly. The sex-dependent evolutionary characteristics of the keystone bacteria strongly correlated with their role played in the microbial community of each sex, i.e. as a keystone for one sex but not for the other, suggesting a role of the evolutionary forces in the shaping of the keystones, which may have further contributed to the formation of the communities. For instance, the nasopharyngeal commensal *N. sicca_D*, acts as the most influential keystone in females while undergoing positive genetic adaptation in response to niche-specific stress condition as of iron limitation, which might have contributed to the formation of the more stable nasal microbial communities against infections. On the other hand, *S. maltophilia_L*, a male-prone respiratory infection associated multidrug-resistant organism, acts as the most influential keystone in males and exhibited strong positive selection for antibiotic resistance relevant efflux pumps, which may further predispose males more vulnerable to infections.

In conclusion, we leveraged in this work the most advanced techniques in the microbiome research field, and applied deep shotgun whole metagenome sequencing, de novo assembly, and network analyses to explore the understudied human nasal microbiome in the largest cohort as of today. Based on that, we constructed a non-redundant nasal bacterial MAGs catalog, and revealed extensive sex differences in the nasal microbiome of healthy young adults. The results provide valuable insights into the observed discrepancies between males and females in respiratory tract diseases, and will help further our understanding of the microbial roles in pathology and etiology. Nevertheless, the findings are limited to mathematical modeling and inference, and experimental validation is desired in the future. Besides, interactions between viruses and bacteria widely exist, such as the synergism between influenza virus and *S. pneumoniae* (Bosch et al., 2013; Korten et al., 2019; McCullers, 2006). Though females are less contracted with most types of RTIs, they are indeed more vulnerable to certain respiratory viral pathogens, such as influenza (Klein et al., 2012). While we are in short of reliably profiled virome data, antagonistic potentials against influenza as well as other specific pathogens require further investigation.

## Author contributions

R.G., T.Z., and H.J. conceived and directed the project. J.W. initiated the overall health project. H.Y., X.X., X.J., Y.C., P.L., Y.H. and L.X. contributed to the organization of the cohort, the sample collection, and questionnaire collection. W.L., X.T. and Z.J. helped checking the phenotypes. H.L. led the DNA extraction and sequencing. R.G. led the bioinformatic analyses. Y.J., M.L., S.L., and Z.S. performed the bioinformatic analyses, and prepared figures and texts for manuscript. Z.Z. and R.G. interpreted the results. Z.Z., R.G. and Y.J. wrote the manuscript. Q.S. contributed to the revision of the manuscript. All authors read and approved the final manuscript.

## Acknowledgments

We sincerely thank the support provided by the China National Gene Bank. We thank all the volunteers for their time and contribution. The data that support the findings of this study have been deposited into CNGB Sequence Archive (CNSA)(Guo et al., 2020) of China National GeneBank DataBase (CNGBdb)(F. Z. Chen et al., 2020) with accession number CNP0002487.

## Data and code availability

Metagenomic data have been deposited into the Genome Sequence Archive (GSA) with accession number CRA006819 and CNSA of CNGBdb with accession number CNP0002487.

## Methods

### Collection of the nasal microbiome samples

Extensive metadata and different biological samples were collected during physical examination in the 4D-SZ cohort as previously reported (Zhu et al., 2021). In this study, we collected anterior nare swabs from 1593 individuals of this cohort, with an average age of 29.9 (±5.13) years old, and sex information obtained for 439 males and 807 females. Demographic characteristics of the participants were provided in Table S1.

The anterior nare samples were self-collected by the volunteers following three steps. First, the sterile swab was moistened with sterile water before use. Then the pre-moistened swab rotated three times around the inside of each nostril with approximately constant pressure. Last, dropping the swab into the 2ml BGI stabilizing reagent (Han et al., 2018) for the preservation of metagenome at room temperature and then stored at -80°C for long-term storage.

### DNA extraction, sequencing, and quality control

DNA extraction of the stored samples was performed using the MagPure Stool DNA KF Kit B (MD5115, Magen) (Yang et al., 2020). Metagenomic sequencing was performed on the DNBSEQ platform (BGI, Shenzhen, China) (Q. Li et al., 2019) with 150 bp of paired-end reads, which generated 854.7 billion pairs of raw reads (on average 536.5 million paired reads per sample, 159.6 million pairs of standard deviation). The metapi pipeline (https://github.com/ohmeta/metapi) was used to process the sequencing data. Quality control was first performed with strict standards for filtering and trimming the reads (average Phred quality score ≥ 20 and length ≥ 30) using fastp v0.20.1 (S. Chen et al., 2018). Human reads were then removed using Bowtie2 2.4.2 (Langmead and Salzberg, 2012) (human genome GRCh38). In total, 4.2 terabases of high-quality paired-end reads were retained with average 96.35% host ratio (Table S2).

### Recovery of the bacterial community

A single sample assembly and single sample binning strategy was employed to reconstruct bacterial genomes from the preprocessed data using the metapi pipeline. Specifically, the high-quality reads of each sample were individually assembled by applying MEGAHIT v1.2.9 (D. Li et al., 2015) or SPAdes v3.15.2 (Nurk et al., 2017) (--meta). BWA-MEM v0.7.17 (H. Li and Durbin, 2009) with default parameters was then used to map reads back to the contigs, and the contig depth was calculated by jgi_summarize_bam_contig_depths (Kang et al., 2019). Metagenomic binning was performed with DAS Tool 1.1.2 (Sieber et al., 2018), combining CONCOCT v1.1.0

(Alneberg et al., 2014), MaxBin v2.2.7 (Y.-W. Wu et al., 2016) and MetaBAT2 v 2.15

(Kang et al., 2019) for each sample individually. CheckM v1.1.3 (Parks et al., 2015) was used to assess the quality of the MAGs. Bins with ≥ 80% completeness and ≤ 10% contamination were retained for further analysis (Stewart et al., 2018). All of the MAGs were then together dereplicated by dRep v3.0.1 (-pa 0.9 -sa 0.99 -nc 0.30 -cm larger -p 25)

(Olm et al., 2017), in which the primary cluster using MASH with 90% ANI and the secondary cluster using ANImf with 99% ANI, resulting in 974 non-redundant MAGs. The 16S rRNA sequences in the MAGs were searched by Barrnap v0.9 (--reject 0.01 --evalue 1e-3, https://github.com/tseemann/barrnap) and tRNA sequences in the MAGs were searched by tRNAscan-SE 2.0.7 (Chan and Lowe, 2019) with default parameters. Taxonomic classification of the 974 non-redundant MAGs was assigned using GTDB-Tk v1.5.1 (Chaumeil et al., 2019) classify workflow with external Genome Taxonomy Database release 95. The phylogenetic tree of the 974 MAGs was built using GTDB-Tk v1.5.1. Genome-wide functional annotation was performed using EggNOG mapper v2.1.3 (Cantalapiedra et al., 2021) based on EggNOG v5.0 database (Huerta-Cepas et al., 2019). The bacterial biome profile was then generated using CoverM with genome mode (--min-covered-fraction 0) (https://github.com/wwood/CoverM) based on the non-redundant nasal bacterial MAGs catalog.

### Characterization of fungal community composition

High-quality cleaned reads were mapped to a manually curated database using kraken2 with default parameters to generate the fungal biome profile. This database contained 39,559 species in total, including human genome GRCh38, GTDB r95, fungi and protists from NCBI.

### Unsupervised clustering

The weighted similarity network fusion (WSNF) analysis (Mac Aogáin et al., 2021) can integrate multi-biome data and cluster samples into distinct groups using taxonomic richness of each biome as the weight of SNF. In this study, for 1593 participants, we filtered bacteria and fungi with relative abundance greater than 1e-4 and 1e-3, respectively, in addition to a prevalence greater than 10%. Finally, 122 bacteria and 131 fungi remained. Three clusters were derived with WSNF from the filtered dataset. Other parameters are set as default.

### Co-occurrence analysis of microbial interaction

To mitigate the influence of spurious correlation, a modified co-occurrence analysis based on ensemble methods was implemented (Faust et al., 2012). In this study, we made modification of this co-occurrence analysis by replacing some methods in the ensemble. First, we implemented COAT (composition-adjusted thresholding) (Cao et al., 2018) instead of Spearman and Person correlation. Then we replaced HUGE (High-dimensional Undirected Graph Estimation) (Zhao et al., 2012) with GBLM (generalized boosted linear models). Last, the sign of the correlation depends on COAT and HUGE. Overall, the ensemble contained MI (mutual information), Bray-Curtis dissimilarity, COAT and HUGE. The final interaction score aggregated the normalized absolute edge scores, and the sign was assigned based on COAT and HUGE. The final *P*-value was merged using the weighted Simes test.

With the modified ensemble method, we conducted co-occurrence analysis on the filtered nasal microbiome dataset (as described in **Unsupervised clustering)** for males and females. Filtering out of the low abundance and low prevalence taxa of the microbiome data helped to avoid artificial interactions resulted from random noises though at the expense of sensitivity loss for weak signals. Following the co-occurrence analysis, the nasal microbial interaction networks were established with a threshold of P-value lower than 1e-3 for males and females.

### Stability of microbial co-occurrence network

Natural connectivity is a robustness measure of complex networks (Morone and Makse, 2015). Higher natural connectivity indicates higher network stability. In this study, we performed random attack by removing randomly selected nodes for 1000 times and assessed normalized natural connectivity for each remaining network (R package pulsar). The number of nodes removed was sequentially increased from 1 to all the nodes. The *P*-value of robustness between male and female networks was calculated following two steps. First, for each attacked network, compare the 1000 natural connectivity between the two sexes with Wilcoxon rank-sum test. Second, a merged P-value was measured using the weighted Simes test, with the number of remaining nodes as the weight.

### Selection of keystone taxa in the co-occurrence networks

We selected keystones based on the IVI (Integrated Value of Influence) (Salavaty et al., 2020), which is an integrative method for the evaluation of node influence within a network. To determine the keystones of each network, we utilized a permutational approach by comparing each robustness attack along the IVI decreasing axis with random attacks (as control). The keystone set was then decided based on *P-*value < 0.001 calculated from the 1000 permutations. Through analysis, we got 13 and 10 key players of male and female network, respectively.

### pN/pS ratios

SNVs of nonsynonymous and synonymous variants at the gene and genome levels were identified for the keystone taxa using inStrain (Olm et al., 2021). The pN/pS ratio was calculated using the formula ((nonsynonymous SNVs/nonsynonymous sites)/(synonymous SNVs/synonymous sites)).

### BGCs prediction

BGCs (biosynthetic gene clusters) type and location of non-redundant MAGs were predicted using AntiSMASH 6.0.0 (Blin et al., 2021) (--cb-knownclusters). Novel BGCs were defined which did not match the Minimum Information about a Biosynthetic Gene cluster (MIBiG) database.

### Statistical analysis and data visualization

#### LDA analysis

For discriminant analysis of the microbiome between males and females, the effect size of linear discriminant analysis was implemented using the webtool available at http://huttenhower.sph.harvard.edu/galaxy/

#### Permanova analysis

ADONIS (permutational multivariate analysis of variance using distance matrices) testing between the observed clusters or sex was performed using R package vegan v2.5-7 with 4999 permutations.

#### Diversity analysis

The nasal microbiome α-diversity (within-sample diversity) was calculated using the Shannon index at the species level (R package vegan). The differences between males and females were assessed with Wilcoxon rank-sum test.

#### Correlation of IVI and relative abundance

The correlation between IVI and relative abundance of the keystone taxa was measured by Spearman’s correlation.

#### Visualization

The co-occurrence network was visualized using Cytoscape 3.9.0. The heatmap of similarity score was drawn by ComplexHeatmap(2.10.0). The boxplot was drawn by ggpubr(2.10.0).

