## Supplementary Figure for "Sex differences in the nasal microbiome of healthy young adults"

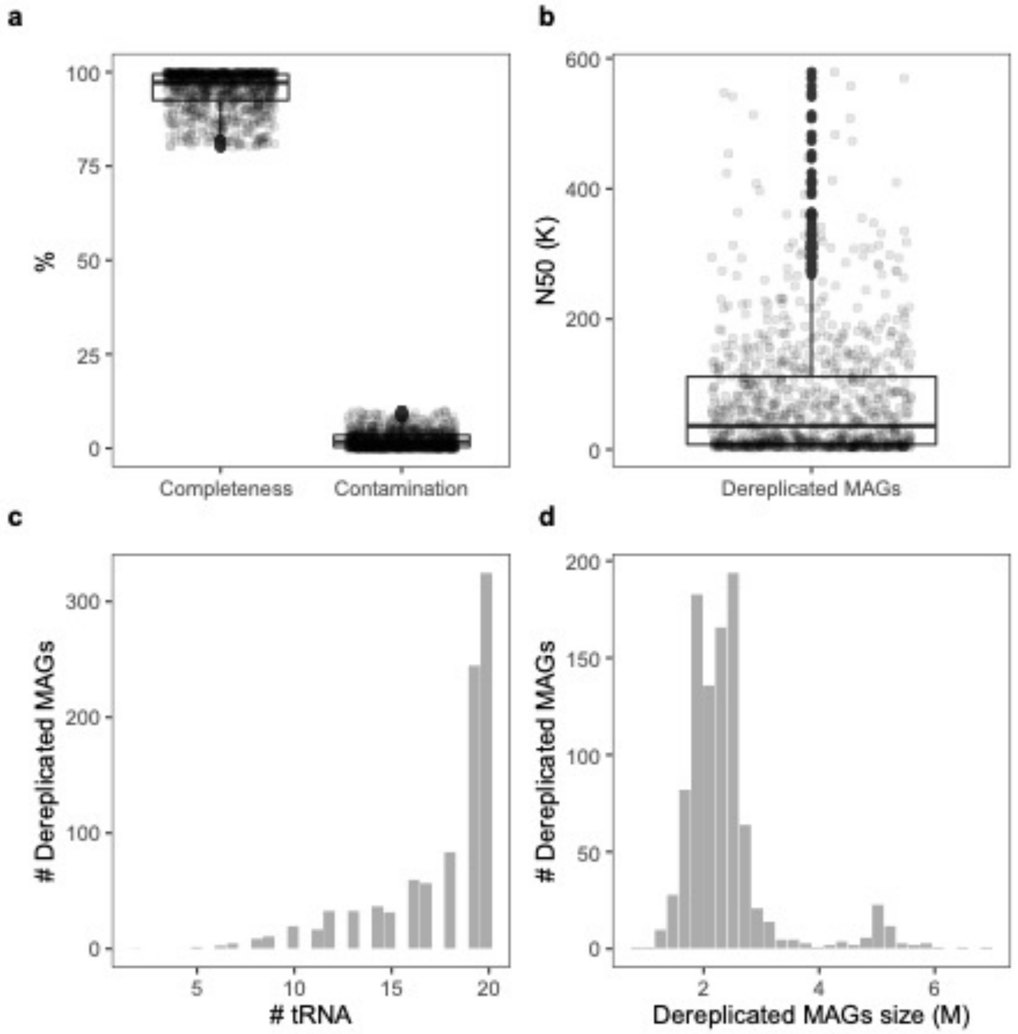


**Supplementary Fig.1 Quality statistics of the 974 non-redundant MAGs.**

**a**, Completeness and contamination level of the 974 non-redundant MAGs. **b**, N50 of the 974 non-redundant MAGs. **c**, Distribution of the number of tRNAs coding for the 20 standard amino acids detected in the 974 non-redundant MAGs. **d**, Distribution of the genome size of the 974 non-redundant MAGs.

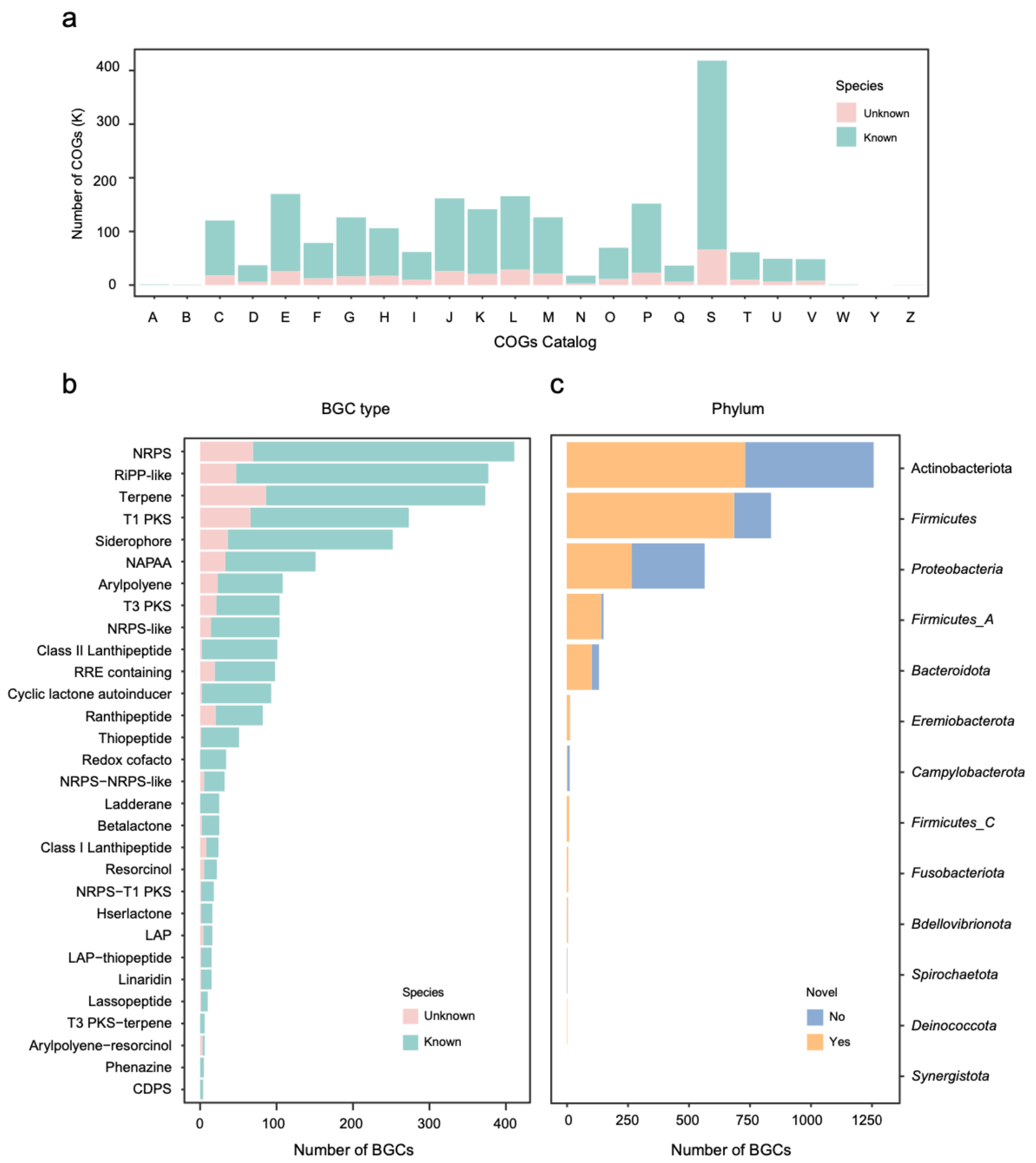


**Supplementary Fig.2 COG functional categories and biosynthetic gene clusters identified in the bacterial species of the nasal microbiome.**

**a-b**, Number of COG functional categories (**a**) and BGCs (**b**) identified in unknown (pink) and known (turquoise) species of the nasal microbiome. **c**, Fraction of all BGCs that could be annotated with the MIBiG database (blue) or not (brown).

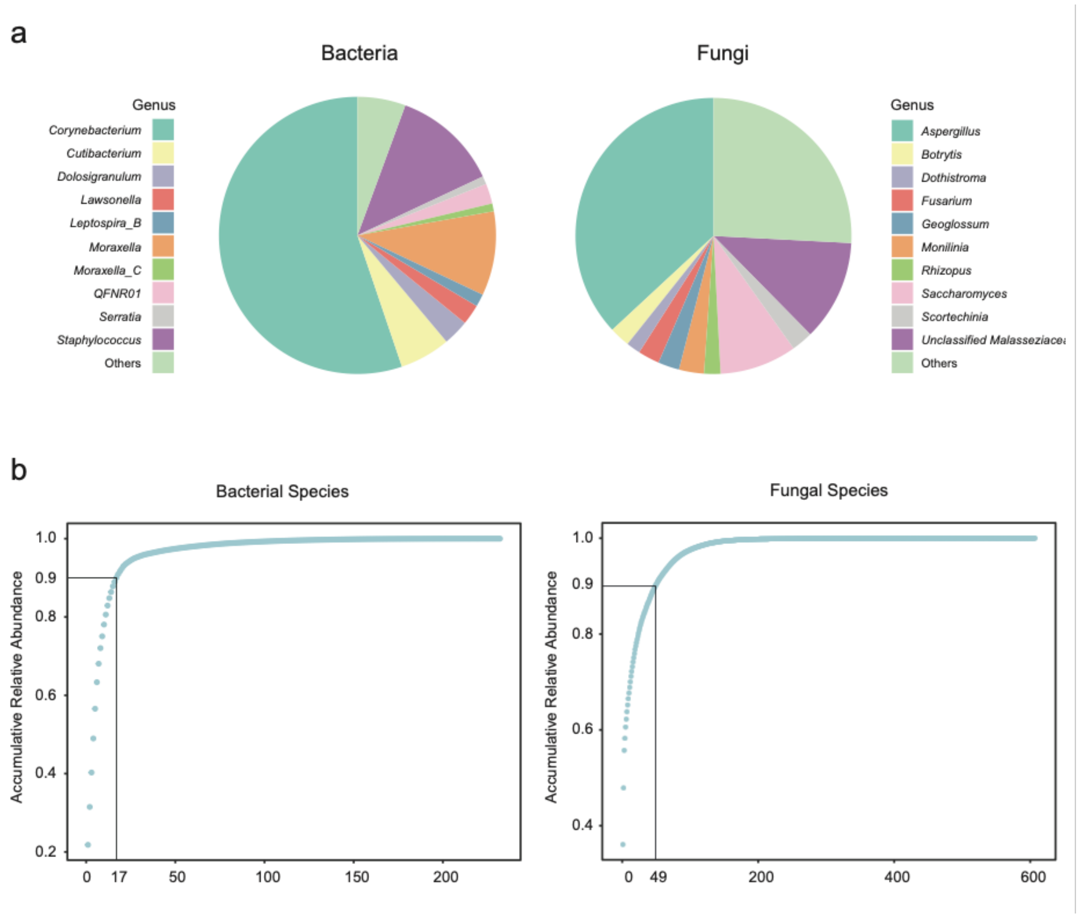


**Supplementary Fig.3 Overall composition of the nasal microbiome**

**a**, Pie charts illustrating the genus level nasal bacterial (left) and fungal (right) biome composition. **b**, Accumulative relative abundance of the nasal bacteriome (left) and mycobiome (right) at the species level.

a

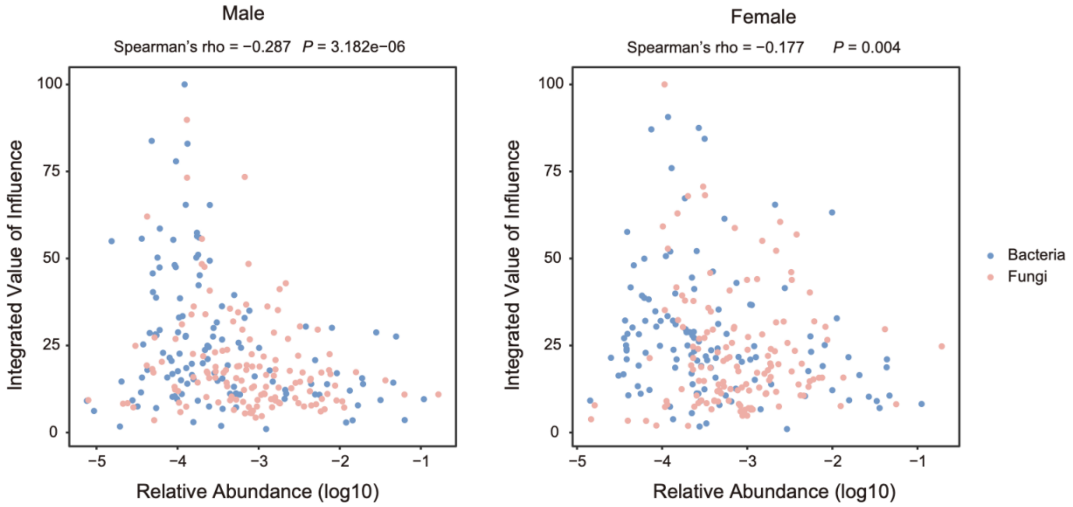


b

| | Bacteria | | Fungi | |
|---|---|---|---|---|
| | Male | Female | Male | Female |
| Spearman's rho | -0.290 | -0.180 | -0.290 | -0.180 |
| *P* | 0.003 | 0.003 | 0.150 | 0.740 |

**Supplementary Fig.4 Correlation between relative abundance and the integrated value of influence in male and female networks for each nasal microbial taxon.**

**a**, scatter plot with Spearman's rho and P calculated on all taxa (bacteria+fungi). **b**, Spearman's rho and P calculated for bacteria and fungi separately.

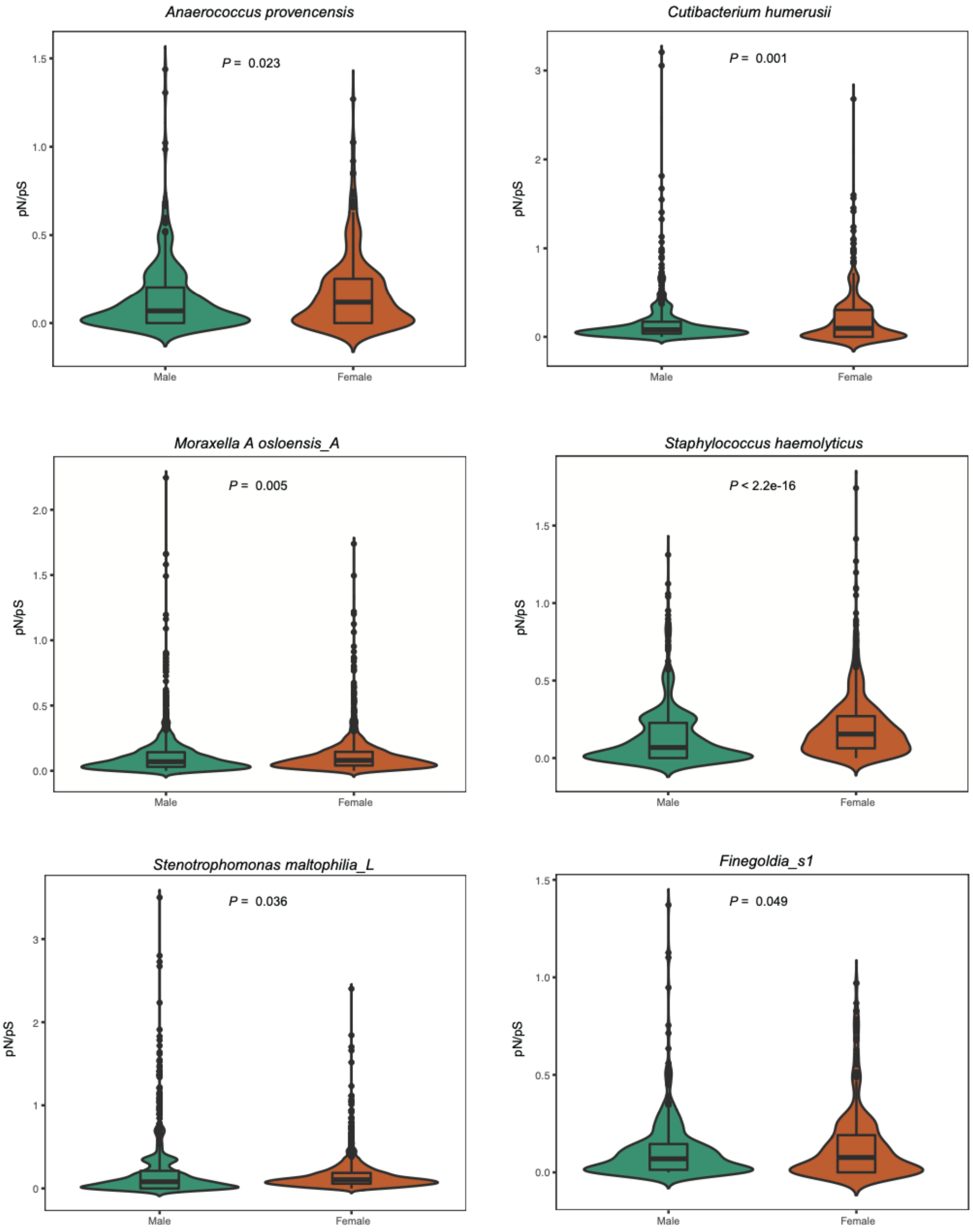


**Supplementary Fig.5 Comparison of pN/pS ratios for genes in keystones between males (green) and females (brown).**

Comparison of pN/pS ratios for genes in keystones between males (green) and females (brown). one-tailed Wilcoxon rank-sum test ($p < 0.05$).
