## Supplementary figures and images for "Sex differences in the nasal microbiome of healthy young adults"

### Figure S1

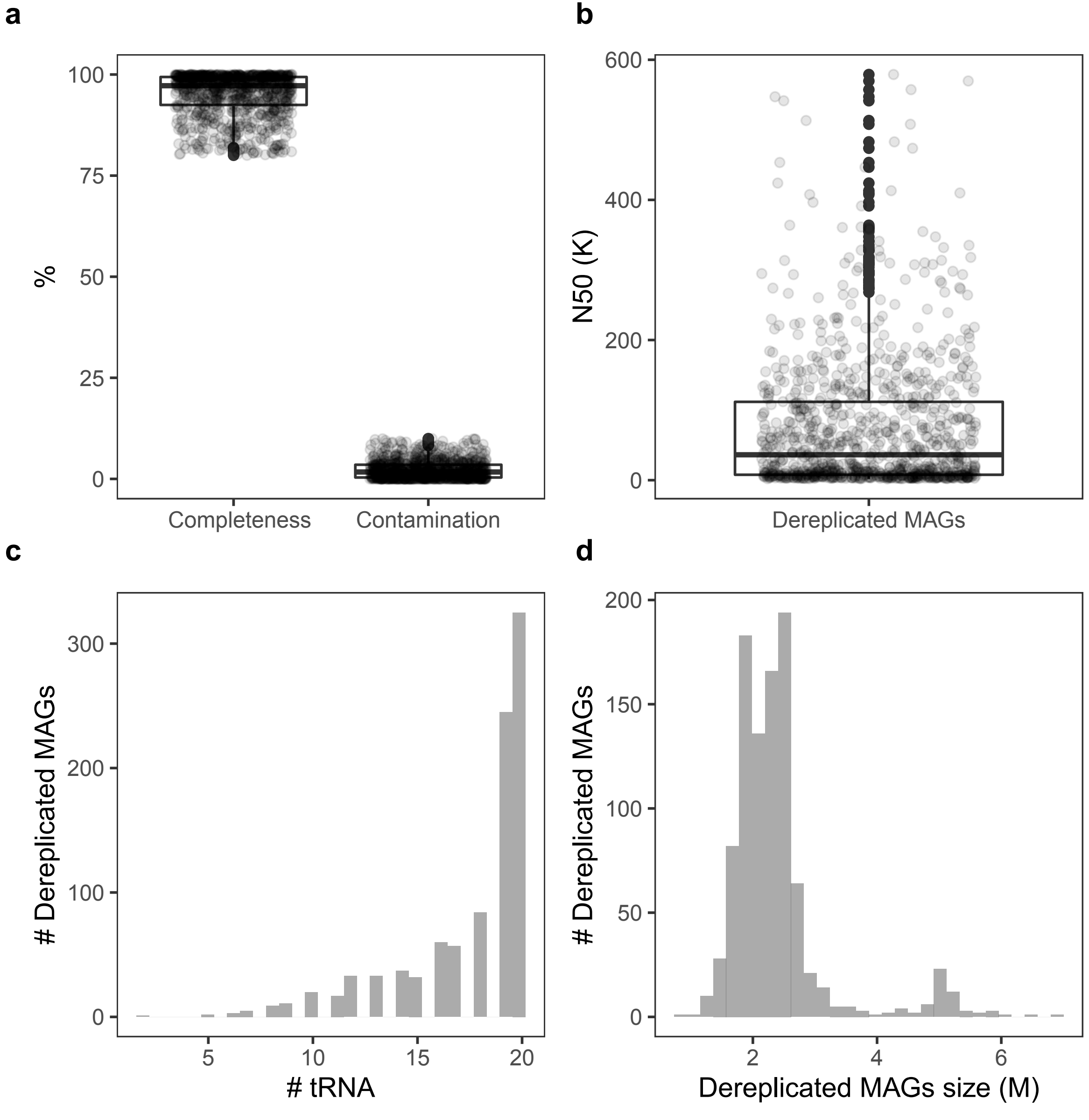

### Figure S2

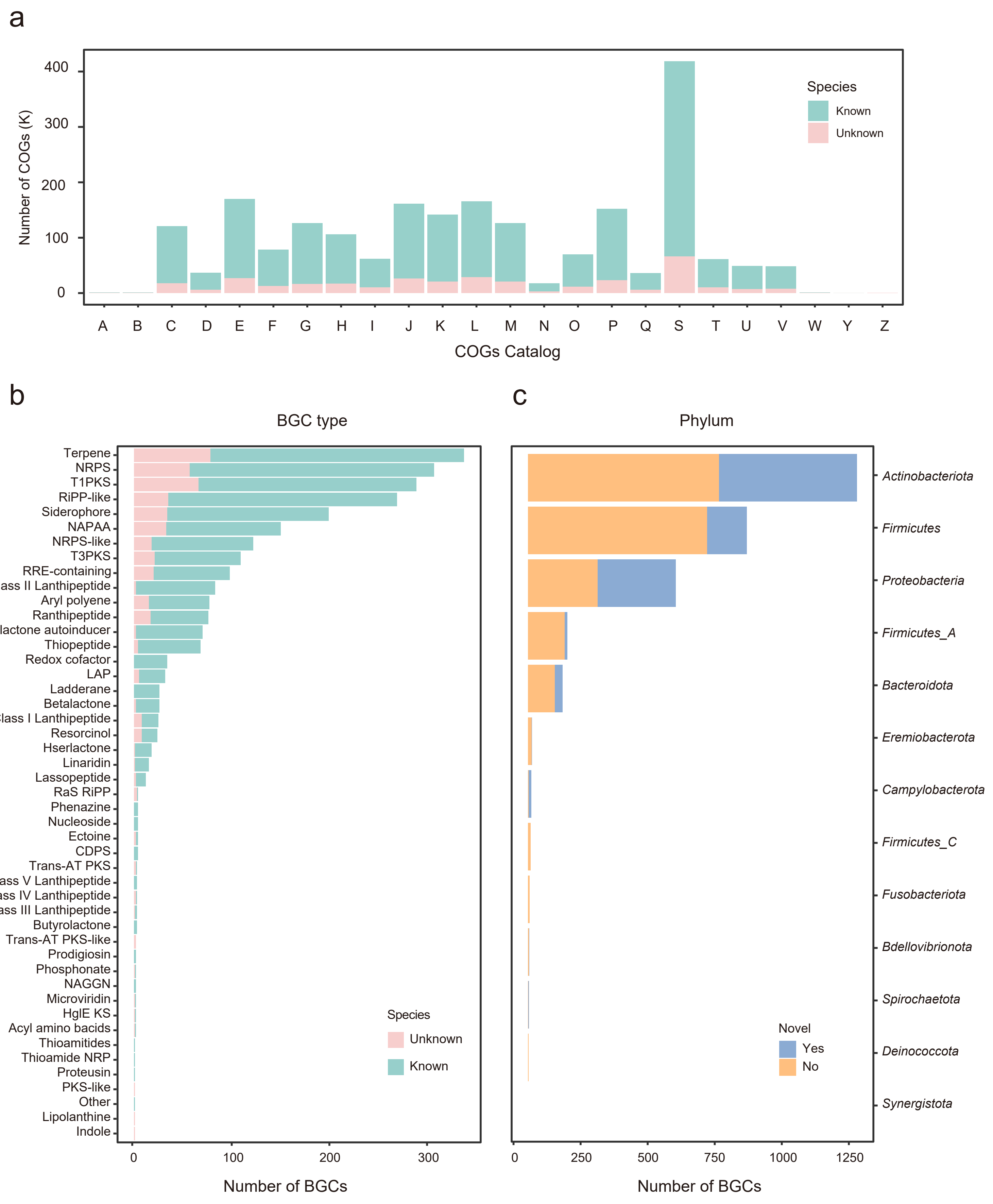

### Figure S3

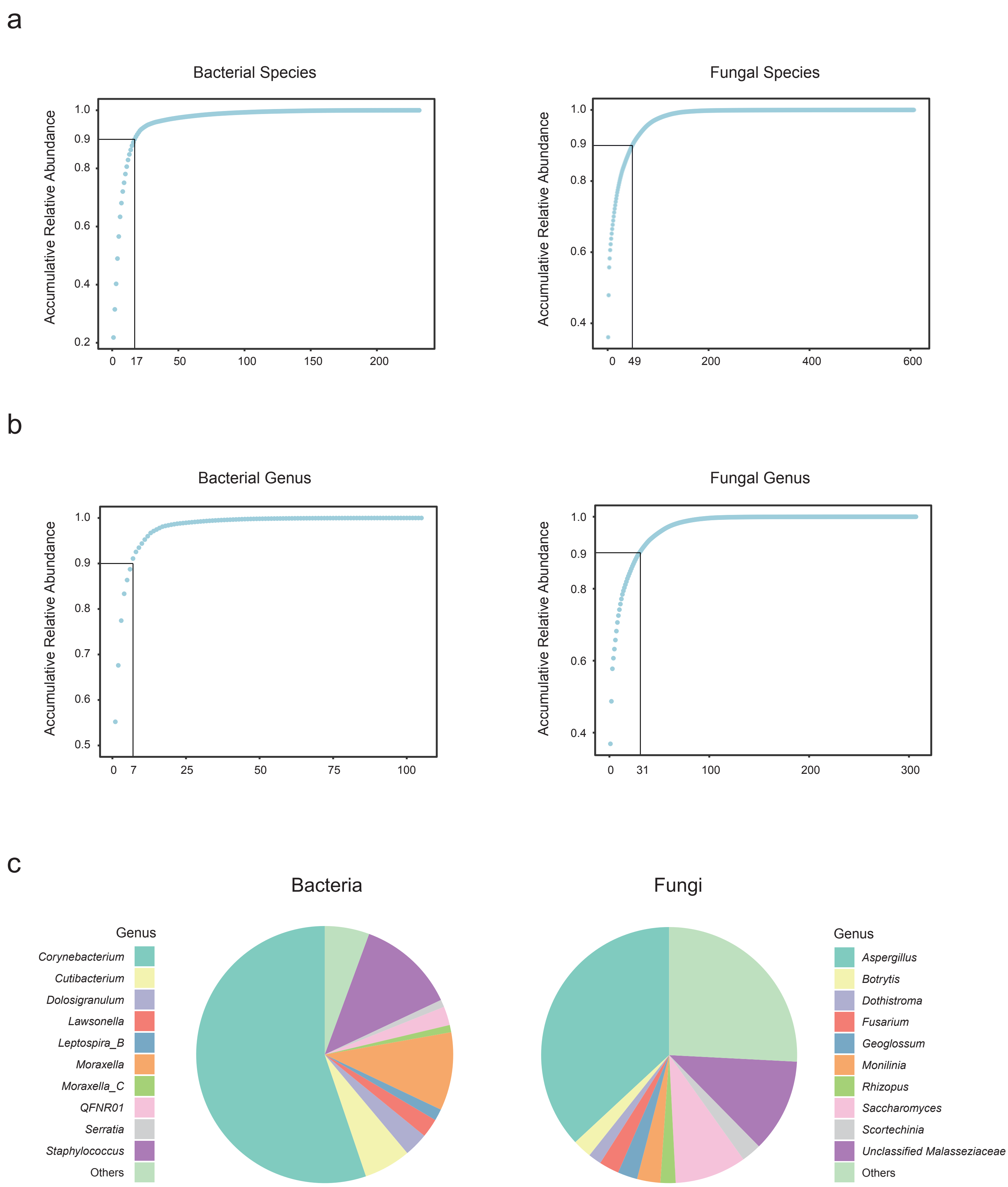

### Figure S4

a

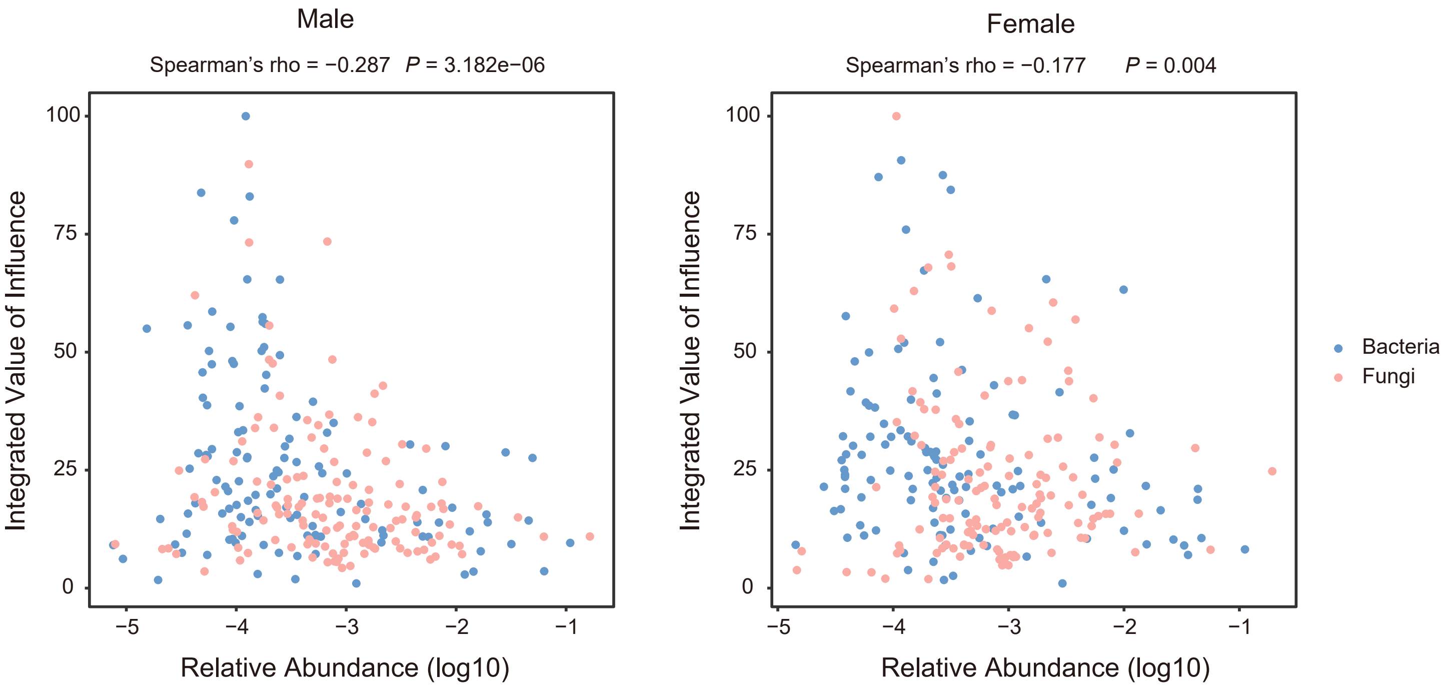


b

| | Bacteria | | Fungi | |
|---|---|---|---|---|
| | Male | Female | Male | Female |
| Spearman's rho | -0.290 | -0.180 | -0.290 | -0.180 |
| *P* | 0.003 | 0.003 | 0.150 | 0.740 |

### Figure S5

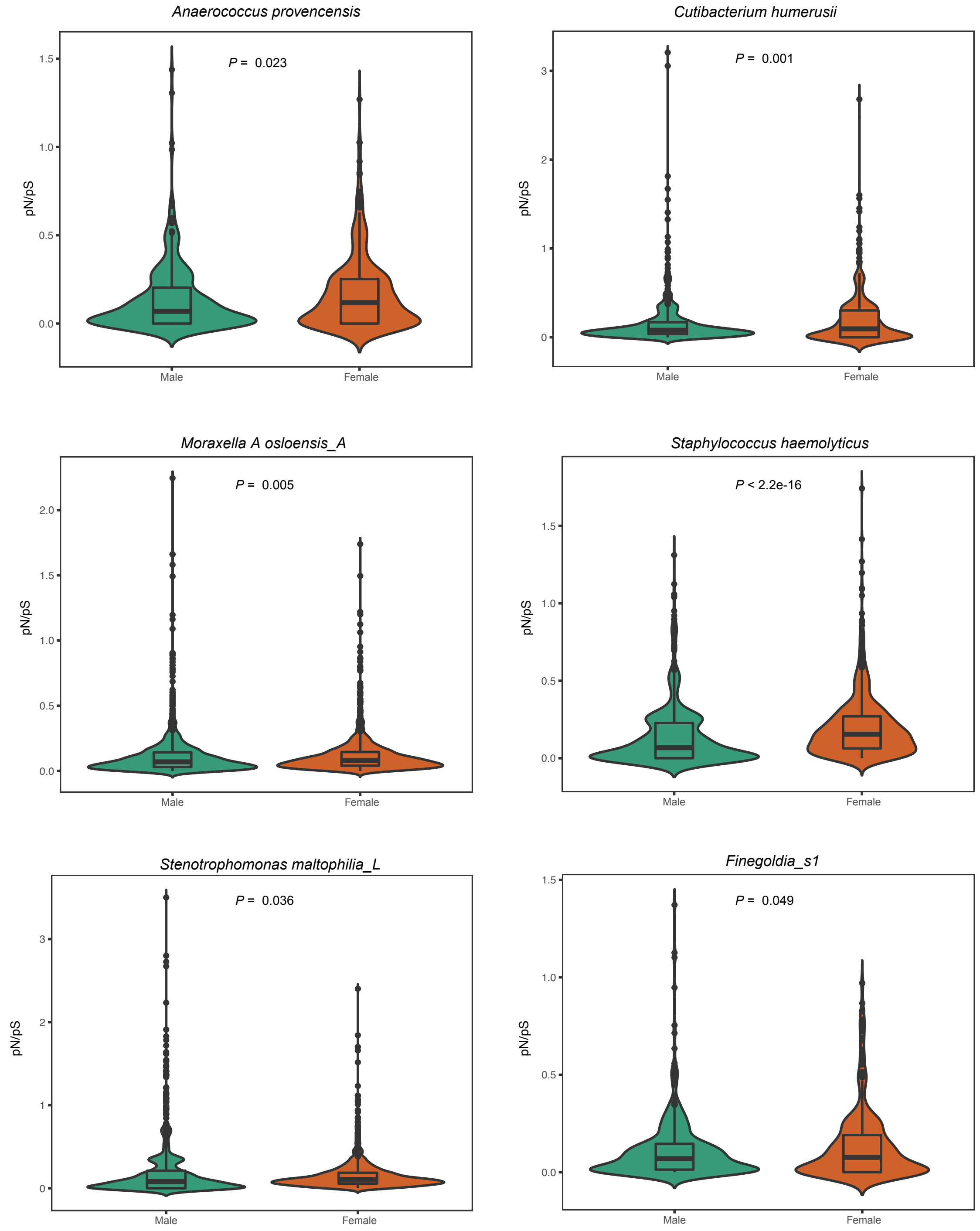
